# ZELDA: a 3D Image Segmentation and Parent-Child relation plugin for microscopy image analysis in *napari*

**DOI:** 10.1101/2021.10.24.465596

**Authors:** Rocco D’Antuono, Giuseppina Pisignano

**Affiliations:** Crick Advanced Light Microscopy STP, The Francis Crick Institute, London, United Kingdom; Department of Biology and Biochemistry, University of Bath, Bath, United Kingdom

**Keywords:** image analysis, 3D, segmentation, Parent-Child, napari, plugin, microscopy, measurement

## Abstract

Bioimage analysis workflows allow the measurement of sample properties such as fluorescence intensity and polarization, cell number, and vesicles distribution, but often require the integration of multiple software tools. Furthermore, it is increasingly appreciated that to overcome the limitations of the 2D-view-based image analysis approaches and to correctly understand and interpret biological processes, a 3D segmentation of microscopy data sets becomes imperative.

Despite the availability of numerous algorithms for the 2D and 3D segmentation, the latter still offers some challenges for the end-users, who often do not have either an extensive knowledge of the existing software or coding skills to link the output of multiple tools. While several commercial packages are available on the market, fewer are the open-source solutions able to execute a complete 3D analysis workflow.

Here we present ZELDA, a new *napari* plugin that easily integrates the cutting-edge solutions offered by python ecosystem, such as *scikit-image* for image segmentation, *matplotlib* for data visualization, and *napari* multi-dimensional image viewer for 3D rendering. This plugin aims to provide interactive and zero-scripting customizable workflows for cell segmentation, vesicles counting, parent-child relation between objects, signal quantification, and results presentation; all included in the same open-source *napari* viewer, and “few clicks away”.

## 1 Introduction

Microscopy and image analysis significantly contribute to the advancement of research in life sciences. However, researchers operating microscopes have to deal with a number of experimental challenges often requiring different types of image analysis procedures. For example, the counting of protein structures, such as the ProMyelocytic Leukemia Nuclear Bodies (PML NB) found involved in chromatin remodeling, telomere biology, senescence or viral infections [Lallemand-Breitenbach, 2018], is achievable by applying a “2D counting” image analysis tool to first identify cells and then determine the number of contained PML NB (**Suppl. Fig. 1A**). Similarly, in experiments where the measurement of transient concentration of Ca2+ or metabolites is assessed, a stable staining and reliable segmentation of individual cytoplasmic organelles might be required to then apply a “2D measurement” of fluorescence intensity and organelle shape (**Suppl. Fig. 1B**). This can be fundamental in studies of mitochondrial metabolism where a complex correlation between ER-mitochondria Ca2+ fluxes and autophagy have been highlighted [Missiroli, 2020]. Furthermore, some kidney pathological conditions, such as the glomerulocystic disease, could originate from topological defects acquired during development [Fiorentino, 2020]. Such conditions can be studied using a staining to identify single cells, glomeruli, and the renal tubular system (**Suppl. Fig. 1C**). The conformational study of a glomerulus, with the assessment of the number of cells, is referred to as “3D cell counting” or “3D object segmentation”. In influenza infection, instead, the released viral genome can be involved in mechanisms such as replication or viral protein transcription and identified by the presence of a negative-sense RNA [Long, 2019]. The dynamics of the viral infection can therefore be monitored by localizing the RNA molecules within the cell nuclei (**Suppl. Fig. 1D**) in a task definable as “3D object segmentation” and “parent-child relation”.

The ability to extrapolate valuable results from microscopy experiments as those just mentioned, mainly relies on the image analysis knowledge and availability of the right software tools for the specific purpose. The bioimage analysis is a combination of multiple informatics tools (referred to as “components”) organized into “workflows” with different levels of complexity [Miura, 2020]. Such components are often available only by scripting and researchers may struggle to find an effective way of combining them together in a complete workflow. To date, there have been great initiatives to both promote the bioimage analysis (NEUBIAS Training Schools [Martins, 2021]) and raise awareness about informatics tools (BioImage Informatics Index, http://biii.eu/), while a growing number of excellent open-source software became available [Schindelin, 2012] [McQuin, 2018]. However, the end-user has still to acquire a minimum level of bioinformatic knowledge in order to analyze image data.

A recent survey proposed by the COBA^1^ to the bioimage analysis community has suggested that the most used bioimage analysis tools belong to the category of the “open-source point and click software” and there is a high demand for better software for “3D/Volume” and “Tissue/Histology” analysis [Jamali, 2021], underlining the urgency of more and more new, easy and customizable tools for multi-dimensional image segmentation.

Furthermore, to guarantee the experimental reproducibility, minimize the mistakes, and preserve scientific integrity, any new analysis software should include accurate logging of the used parameters at each step of the workflow^2^.

To facilitate life science researchers during the application of image analysis to biological experiments, we developed ZELDA: a *napari* plugin for the analysis of 3D datasets with multiple object populations. ZELDA has the advantage of being equipped with ready-to-use protocols for 3D segmentation, measurement, and “parent-child” relation between object classes. It then allows the rapid cell counting, quantification of vesicle distribution, and the fluorescence measurement of subcellular compartments for most biological applications. Since each image analysis workflow is designed as a simple protocol with numbered steps, it requires no knowledge of image analysis and it’s sufficient to follow the step-by-step instructions to perform a complete analysis. Furthermore, while the integration in *napari* allows to easily view each step of the image processing as 2D slice or 3D rendering, the visibility, opacity and blending modulation facilitates the tuning of the used parameters (for example threshold value or gaussian filter size) by visualizing multiple layers at the same time.

Altogether ZELDA plugin is a new easy to use open-source software designed to assist researchers in the most common bioimage analysis applications without requiring any scripting knowledge.

## 2 Materials and Methods

### 2.1 Image acquisition

The data sets of influenza infected human eHAP cells, BPAE cells (Invitrogen FluoCells Slide #1) and mouse kidney tissue (Invitrogen FluoCells Slide #1), shown and analyzed in (**Fig. 2**, **Fig. 4**, **Fig.5**, **Suppl. Fig. 1**, and **Suppl. Fig. 3**) have been acquired with a Zeiss LSM880 confocal microscope, using a Plan-Apochromat 20X/0.8 NA objective. A sequential acquisition for DAPI (excitation 405 nm, detection in the range 420nm - 462 nm), AlexaFluor 488 and AlexaFluor 568 (excitation 561 nm, detection in the range 570nm - 615 nm) was used to acquire z-stacks with the total size up to 13 um, every 0.5 um. Pixel size was 0.20 um.

The beads used to show the segmentation workflow (**Fig. 1**) were TetraSpeck™ Microspheres, 0.1 μm; images were acquired on a Zeiss Observer.Z1 using Micro-Manager (https://micro-manager.org/) software with a Hamamatsu ORCA-spark Digital CMOS camera, using a 63X/1.4 NA objective. Pixel size is 0.08 um.

**Figure 1.**
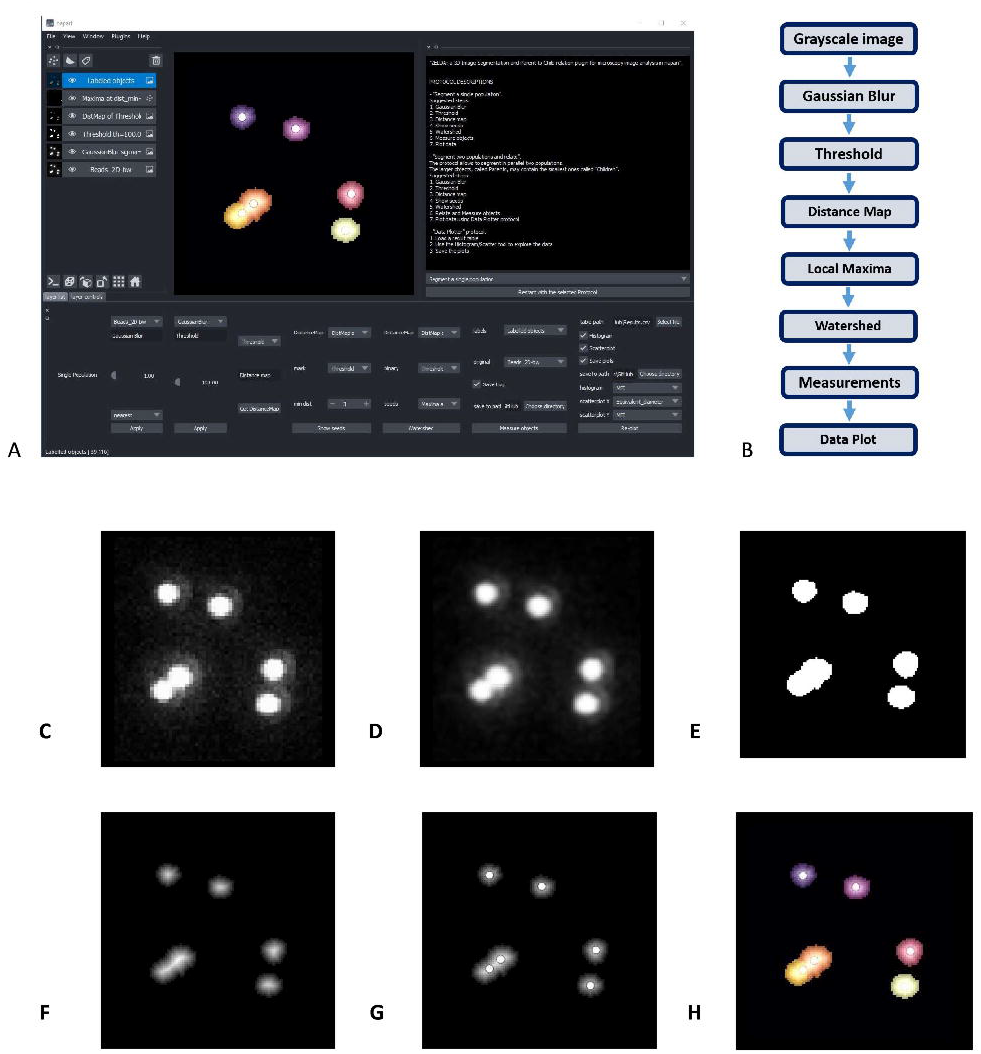
ZELDA plugin for *napari*. (**A**) GUI of ZELDA with the description of the ready-to-use protocols sufficient to run a complete image analysis workflow (white text box on the top right). Each protocol is divided into numbered steps corresponding to the software commands in the *napari* dock widget (bottom of the software interface). (**B**) “Segment a single population” protocol including a minimum number of processing operations. (**C**) Original image. (**D**) Gaussian Blur of the original image. (**E**) Binary image obtained applying a Threshold to the gaussian blur. (**F**) Distance Map applied to the binary image. (**G**) Seeds (Local Maxima) used to run the Watershed. (**H**) Objects labelled by Watershed. The labelled objects can then be measured, and the results exported.

### 2.2 Object segmentation, measurements, and results export

The segmentation obtained by running the ZELDA protocols is achieved using *scikit-image* [van der Walt, 2014] (version 0.18.1) and *SciPy* [Virtanen, 2020] (version 1.6.3) modules for image processing in python.

The resulting measurements are handled as *Pandas* data frames [McKinney, 2011] (version 1.2.4) and plotted with *Matplotlib* [Hunter, 2007] (version 3.4.2).

### 2.3 Graphical User Interface (GUI) design, plugin development, installation, and execution

ZELDA plugin for *napari* (“*napari-zelda*”) can be installed through the “Install/Uninstall Package(s)” menu in *napari* [napari contributors, 2019], and its interface can be added with “Plugins/Add dock widget”.

Alternatively, the installation can be done downloading the repository, navigating to it with the *Anaconda* prompt and using the command “pip install -e.” within the downloaded folder.

The plugin widgets have been created using *magicgui* [https://github.com/napari/magicgui], while the GUI plots included in the “Data Plotter” protocol are obtained with *matplotlib.backends. backend_qt5agg* [https://matplotlib.org/2.2.2/_modules/matplotlib/backends/backend_qt5agg.html].

The template for the plugin has been obtained from *cookiecutter-napari-plugin* [https://github.com/napari/cookiecutter-napari-plugin].

### 2.4 JSON database for modularity of the GUI and customization of image analysis protocols

Once the user has selected a specific base protocol, a JSON file is used by the plugin to load the right widgets in the GUI.

The “Design a new Protocol” option saves the custom workflow as a list of widgets that will be sequentially loaded the next time that the newly created protocol is launched. It will be visible just after relaunching *napari* and ZELDA plugin.

## 3 Results

### 3.1 ZELDA Protocols as an easy way to run image analysis workflows for 2D and 3D segmentation

ZELDA plugin for *napari* (“*napari-zelda*”) makes available to the end-user the segmentation, measurement, and “parent-to-child” relation of two object populations. It ultimately allows to plot the results and explore the data in the same Graphical User Interface (GUI).

The current version of the plugin includes three different “protocols” to ease the image analysis of 3D datasets. Each protocol is a set of individual steps (functions) that return images (as *napari* layers), or results (printed plots in .tiff or tables in .csv format).

The first protocol, called “Segment a single population” (**Fig. 1A**), can be used to segment both the 2D or 3D data sets. The basic workflow of this protocol (**Fig. 1B**) includes simple steps, such as Gaussian Blur, Threshold, and Distance Map, to identify the seed points for the subsequent segmentation of the objects of interest. The user can then set the “min dist” parameter in the “Show seeds” function to improve the accuracy of cell counting, before calling the “Watershed” segmentation (**Fig. 1C-H**). The detected objects can eventually be measured and the results table automatically saved (**Suppl. Fig. 2A**).

Similar workflows have been previously implemented in useful tools such as *MorphoLibJ* [Legland, 2016] and in the latest versions of *CellProfiler* [McQuin, 2018], although with the limitation of being exclusively applied to the 2D image analysis, or lacking an embedded and flexible 3D viewer. In contrast, ZELDA provides an integration of a basic 3D object segmentation workflow with *napari* 3D rendering GUI. Notably, in ZELDA the individual workflow steps are also accessible as single functions that can be optionally used, or fine-tuned individually, without having to restart the entire workflow from scratch.

The second protocol, “Segment two populations and relate” (**Fig. 2A**), implements the segmentation of two populations of objects in parallel, using the same workflow described above, with an additional step that allows establishing the “parent-child” relation between the two object populations (**Fig. 2B-D**).

**Figure 2.**
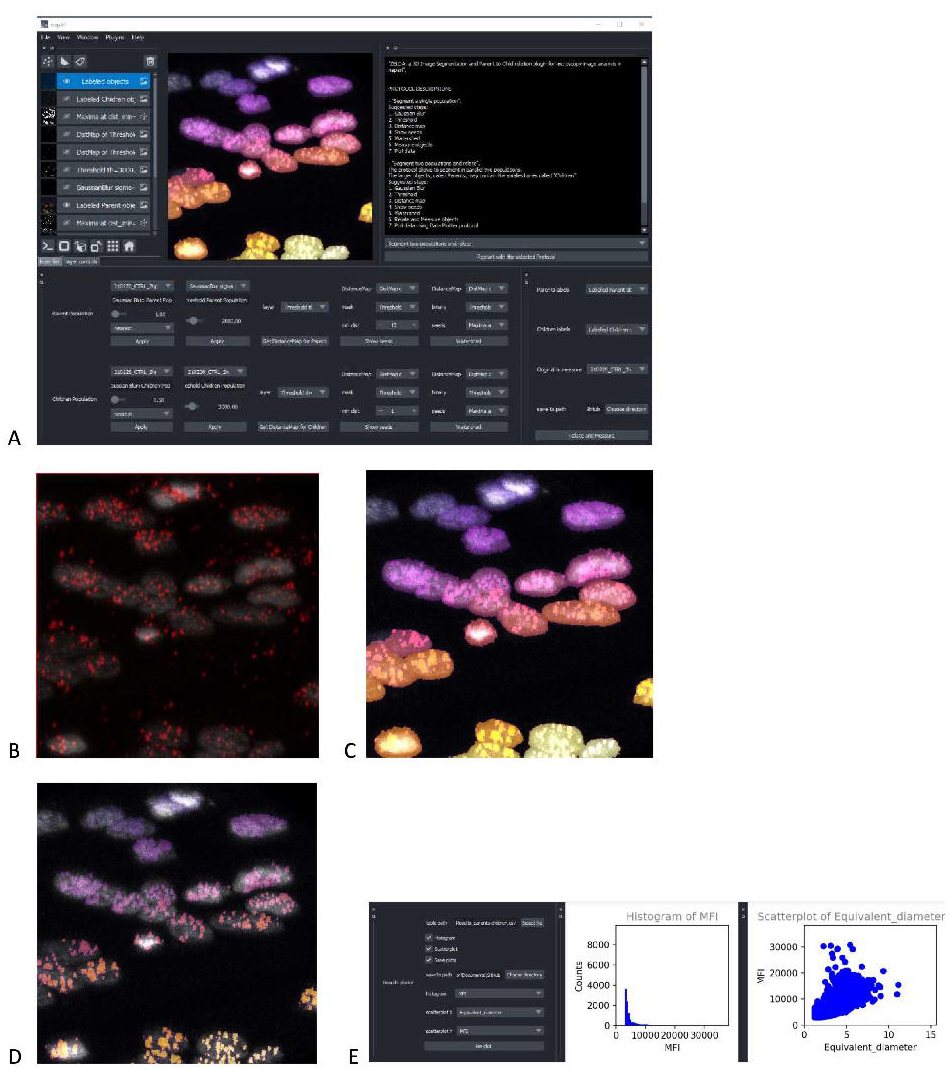
ZELDA application for the 3D segmentation of two object populations and “parent-child” relation. (**A**) ZELDA protocol “Segment two populations and relate” used to analyze the distribution of viral RNA in infected human cell nuclei. (**B**) Original 3D data set showing a nuclear staining with DAPI (gray) and an RNA staining with AlexaFluor 568 (red). (**C**) The nuclei and the RNA aggregates, individually segmented and (**D**) the RNA aggregates (children population) labelled according to the containing nuclei (parent population). (**E**) Resulting measurements reimported and plotted with the “Data Plotter” protocol.

To run reproducible image analysis with ZELDA, both described protocols include a “log” functionality that stores the parameters used at each step. The log is shown in the GUI and can be optionally saved as a .txt file, together with the other results (**Suppl. Fig. 2B**).

Once segmented, measured, and optionally related two object populations, the “Data Plotter” protocol (**Fig. 2E**) allows to load a result table, and plot histograms or scatterplots of the measured properties. The plots are shown directly in the *napari* GUI and can be automatically saved as images to a specific folder. This has the advantage of avoiding the employment of additional software for data visualization.

Given that ZELDA does not require any coding skill, life science researchers are hugely facilitated by the integration of multiple bioinformatics tools in a single GUI.

### 3.2 Modularity of the ZELDA Graphical User Interface allows to easily customize bioimage analysis workflows without any scripting knowledge

Computer scientists and developers continuously propose new algorithms to tackle biological problems that frequently require extensive coding skills. However, users might have the necessity to reproduce a specific published workflow (such as the one in **Fig. 1B**), without knowing a scripting language or necessarily having any background in image analysis. We made this possible by implementing a method that allows the customization of the image analysis protocols available in ZELDA. Indeed, by simply running the fourth option called “Design a New Protocol”, a user can easily create a new custom protocol (**Fig. 3A**). Every step of the base protocols is listed in a JSON database and the relative GUI widgets (used for the software layout) are available as ready-to-use modules to build personalized protocols. The different functions, such as threshold, gaussian blur or distance map etc., can be chosen in a drop-down menu at specific steps of the new protocol (**Fig. 3B**). By using the saving option (**Fig. 3C**), the JSON database will be automatically updated (**Fig. 3D**), and the ordered series of GUI widgets will be available the next time that ZELDA plugin will be launched (**Fig. 3E**).

**Figure 3.**
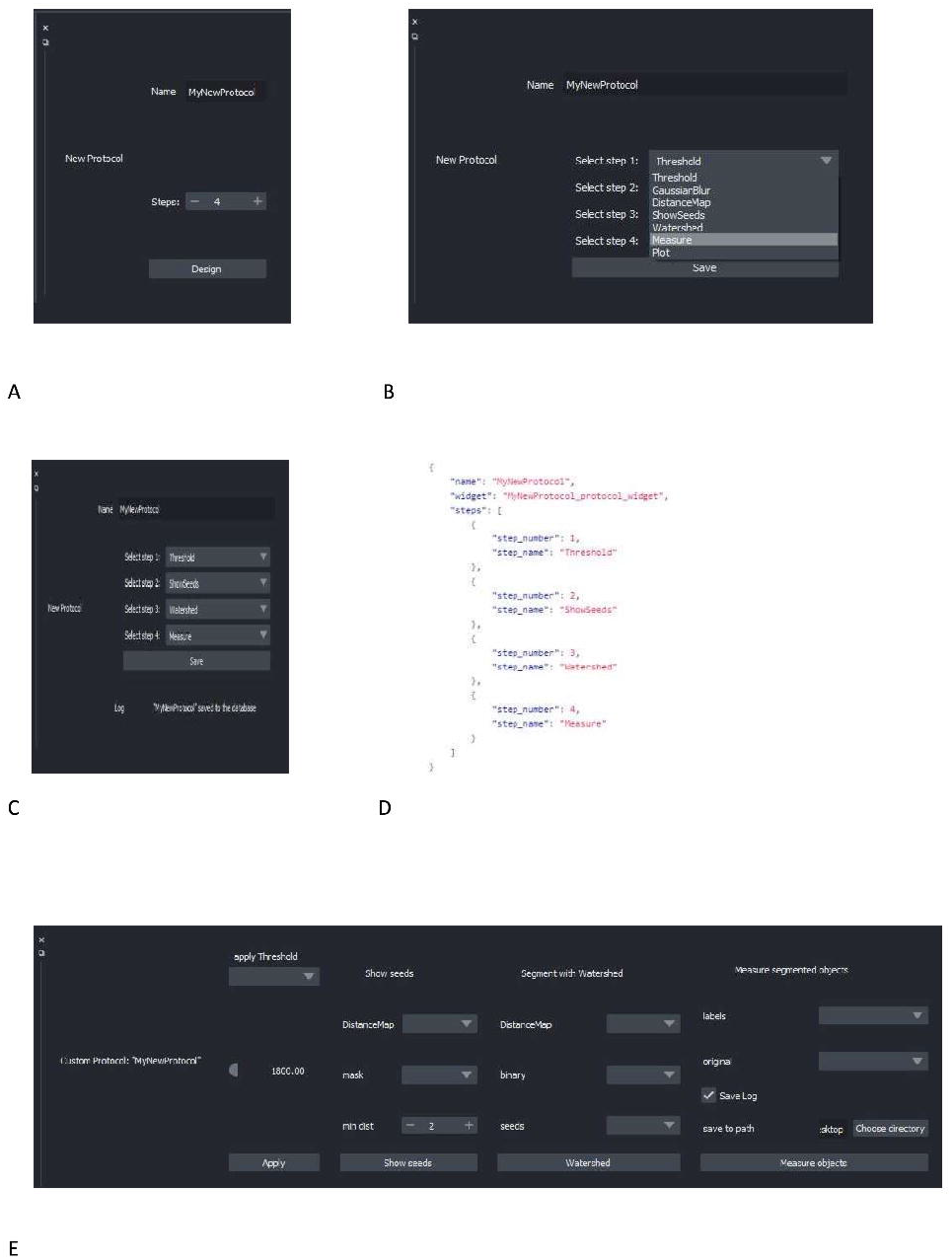
Design of a custom image analysis workflow with ZELDA without requiring any scripting knowledge. (**A**) Choice of the number of steps for the new protocol to implement a custom image analysis workflow. (**B**) Drop-down menu showing all the modules implemented in ZELDA. (**C**) Assignment of an operation to a specific step of the new protocol. (**D**) Example of updated JSON database that controls the software layout, once a new protocol is saved. (**E**) The newly created protocol GUI available after having restarted ZELDA.

### 3.3 ZELDA segmentation and parent-child relation have the same accuracy of *ImageJ* and *CellProfiler* in 2D and 3D data sets, and the execution is twice faster

In order to assess the accuracy in the segmentation of 2D and 3D data sets, we compared the results obtained by the ZELDA plugin for *napari* with those generated by two of the most widely used software in the bioimage field: *ImageJ* and *CellProfiler*.

As 2D data sets, we used images of cells (**Fig. 4A**) at a low confluence (~30% of the field of view area) with a cytoplasmic staining to identify parent objects, and a second one for cellular organelles (children objects), with the final goal of correctly assign the organelles to the containing cell (parent-child relation).

**Figure 4.**
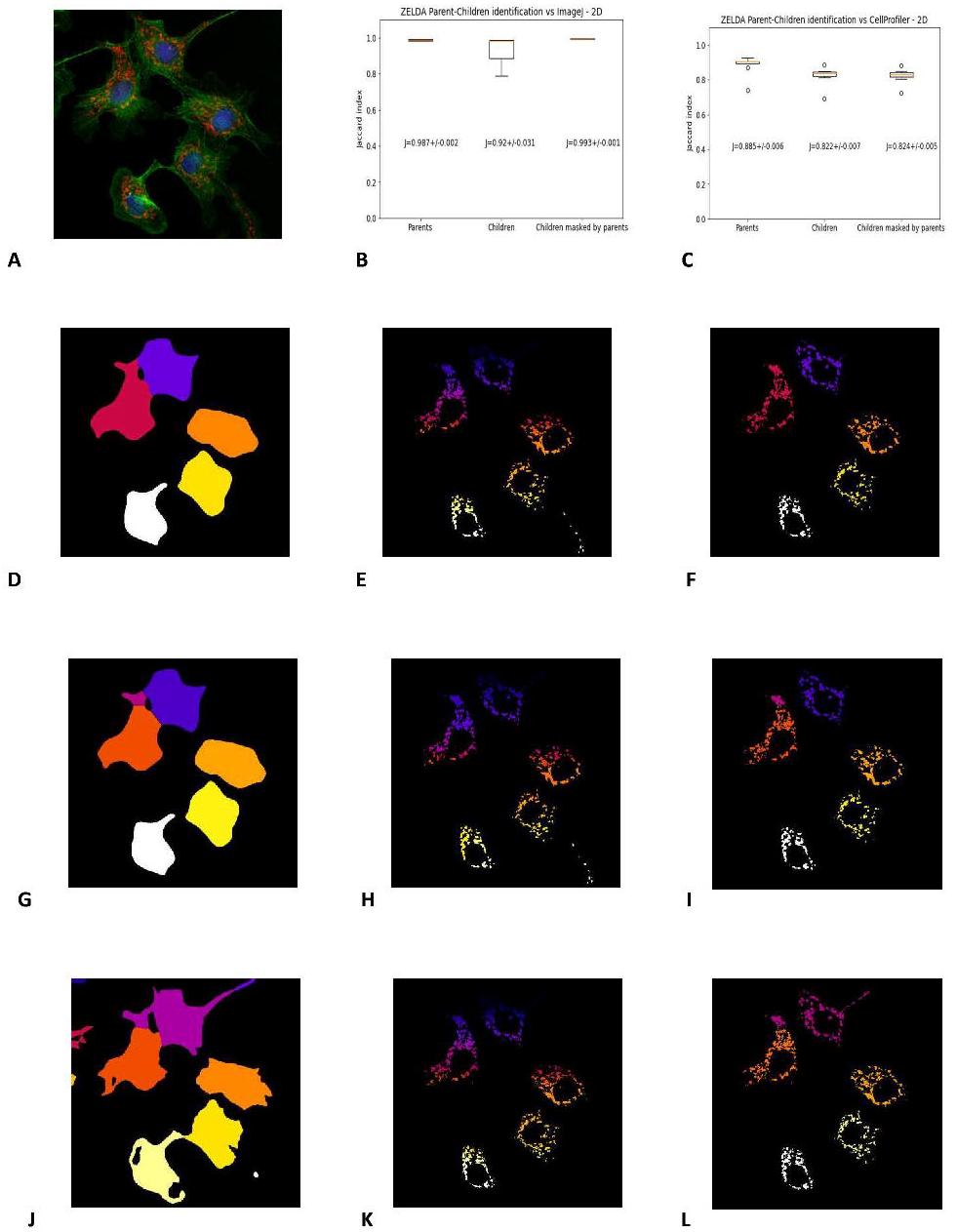
ZELDA 2D segmentation and “parent-child” relation benchmarked with *ImageJ* and *CellProfiler*. (**A**) 2D images of BPAE cells stained with DAPI (blue, cell nuclei), AlexaFluor 488 (green, cytoplasms), and MitoTracker Red (red, mitochondria). (**B**) Labelling comparison between ZELDA and *ImageJ* showing an accordance above the 98 % of the pixels for “parent” objects (cell cytoplasms), 92% for “child” objects (mitochondria), and 99% for the parent-child relation. (**C**) Comparison between ZELDA and *CellProfiler* showing a minor accordance still above the 88% of the pixels for “parents”, 82% for “children”, and 82% for the “parent-child” relation. ZELDA labelling of (**D**) cell cytoplasms, (**E**) mitochondria, and (**F**) masked mitochondria (parent-child relation). *ImageJ* labelling of (**G**) cell cytoplasms, (**H**) mitochondria, and (**I**) masked mitochondria (parent-child relation). *CellProfiler* labelling of (**J**) Cell cytoplasms, (**K**) mitochondria, and (**L**) masked mitochondria (parent-child relation).

Intriguingly, ZELDA performed almost equivalently to *ImageJ* (**Fig. 4B**) in identifying parent objects (Jaccard index J=0.987 +/- 0.002), child objects (J=0.920 +/- 0.031), and in the parent-child relation (J=0.993 +/- 0.001). This means that, assuming *ImageJ* segmentation as ground truth (**Fig. 4G-I**), ZELDA will correctly label the pixels of an organelle as belonging to the corresponding cell cytoplasm in 99% of the cases (**Fig. 4D-F**).

However, the adherence with *CellProfiler* labelling was slightly less striking (**Fig. 4C**) although this difference might be due to the many more parameters available in the *CellProfiler* GUI, such as the “declump method” in the “watershed” module et c., that have not been implemented in ZELDA GUI to keep the software interface and its utilization as simple as possible. Nonetheless, the agreement on the identification of the parent cytoplasms found with *CellProfiler* (**Fig. 4J**) was around 88% of the pixels, while for both the child objects segmentation and the parent-child relation (**Fig. 4K-L**) it was ~ 82%.

Benchmarking the segmentation of 3D data sets has proven to be slightly more complicated, since not all the available modules in *CellProfiler* support the 3D data processing. For example, in version 4.2.1 the “smooth” module that operates a Gaussian blur filter, is available just for the 2D data pipeline, while another one has to be used for the 3D case. The same holds for morphological operations such as those executed by the “ExpandOrShrinkObjects”. Trying to circumvent this lack of interchangeable 2D/3D functions could result in a more elaborated and time-consuming construction of the *CellProfiler* pipeline. Conversely, the versatile protocols supplied with ZELDA (**Fig. 1** and **Fig. 2A-D**) allowed the 3D segmentation and parent-child relation in fewer steps and about twice quicker than the *CellProfiler* “Test mode” (**Fig. 5J**).

**Figure 5.**
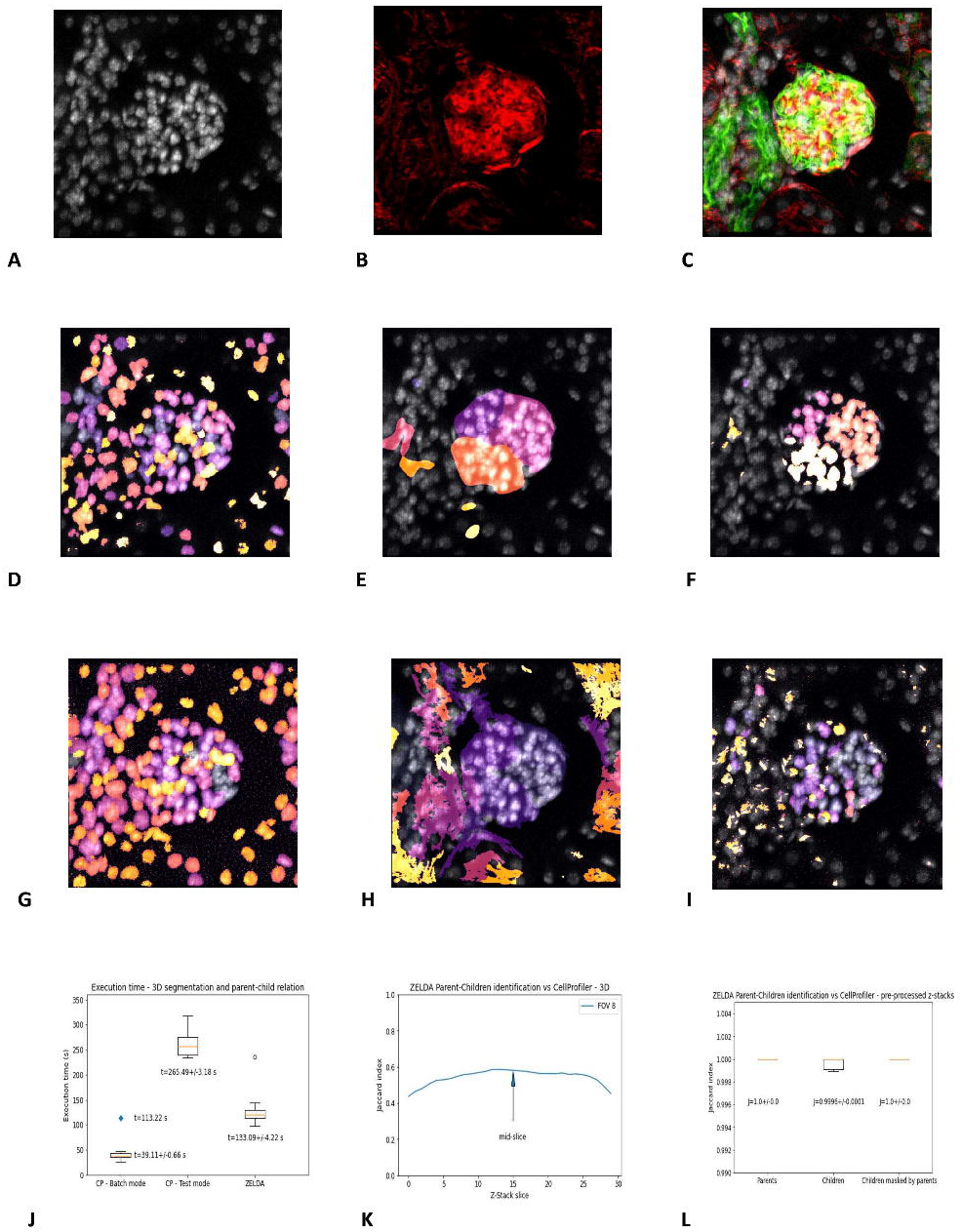
3D segmentation, “parent-child” relation, and execution time of ZELDA compared to *CellProfiler*. Z-stacks of mouse kidney tissue showing glomeruli were used for the benchmark in 3D. (**A**) DAPI staining used to segment the cell nuclei. (**B**) Phalloidin used to identify the glomerular structures. (**C**) 3D rendering showing the merge of DAPI (gray), WGA (green), and phalloidin (red). (**D**) Nuclei, (**E**) glomerular structures, and (**F**) masked nuclei (parent-child relation) labelled by ZELDA in 3D. (**G**) Nuclei, (**H**) glomerular structures, and (**I**) masked nuclei (parent-child relation) labelled by *CellProfiler* in 3D (using a pipeline containing only 3D data compatible modules). (**J**) Execution time for the same workflow developed as both *CellProfiler* pipeline and ZELDA protocol, with the goal to segment in 3D and relate parents and children objects. The boxplots represent the distribution of multiple runs analyzing individual FOVs. For *CellProfiler* in batch mode, the CPU time has been considered, while the blue dot represents the total duration experienced by the end user for the analysis of 9 FOVs (including the wall time). (**K**) Variation of the Jaccard index of the segmentation obtained with ZELDA and *CellProfiler*, around the mid-slice where the signal is stronger. In the 3D case, the maxima of the Jaccard scores along the Z-stack were used for the benchmarking. Not all the *CellProfiler* modules are 3D compatible, then the execution of a minimal pipeline may result in over-segmented structures. The reason was identified to be the lack of a unique name for the same operation in 2D and 3D (“Smooth”), or the absence of 3D equivalents for some modules like the “ExpandOrShrink” morphological operations. *CellProfiler* might be able to process the data sets equivalently to ZELDA but with a longer and more complicated pipeline. (**L**) Increment of agreement on segmentation above the 99% once a pre-processed 3D data by ZELDA was proposed to *CellProfiler*, showing how quickly ZELDA can segment and relate in 3D using less steps than a *CellProfiler* pipeline.

We then analyzed a collection of z-stacks of mouse kidney glomeruli, as 3D data sets (**Fig. 5A-C**). In this tissue, phalloidin staining (**Fig. 5B**) was used for the identification of the glomerular structures, and DAPI staining (**Fig. 5A**) to pinpoint the cell nuclei contained in each glomerulus. The resulting segmentation of the two populations and parent-child relation obtained by ZELDA (**Fig. 5D-F**) were compared with the output of a *CellProfiler* pipeline which included solely the 3D data compatible modules (**Fig. 5G-I**).

Unfortunately, the labelling agreement between the two software was reduced with respect to the 2D analysis. A performance comparison of the 3D segmentation revealed a variation of the Jaccard index across the z-stack, with maximum values typically around the mid-slice, where the staining intensity of the confocal microscopy data set was stronger (**Suppl. Fig. 3A**). We then considered the maxima of the Jaccard index across the z-stacks (**Fig. 5K**), assessing an accordance around 63% for the parent objects (Jaccard index J=0.632 +/- 0.011), 73% for the children (J=0.735 +/- 0.011), and of 64% for the parent-child relation (J=0.643 +/- 0.010) (**Suppl. Fig. 3B**).

We further investigated the reason for the lack of agreement on 3D data sets labelling between ZELDA and *CellProfiler*, and found that the difference was due to the absence of 3D equivalents for some modules (e.g. the “ExpandOrShrink” morphological operations), or lack of a unique naming for the 2D and 3D version of the same method in *CellProfiler* (e.g. “Gaussian Blur”). Indeed, pre-processing the z-stacks with the ZELDA and proposing the resulting smoothed 3D data sets to *CellProfiler*, successfully increased the accordance in identifying parents, children, and parent-child relation above the 99% of the pixels (**Fig. 5L**).

Therefore, ZELDA can represent a faster interactive alternative to *CellProfiler* for the exploratory analysis of 3D data sets.

## 4 Discussion

Many tools are available for 2D segmentation, while fewer are able to process 3D data sets [Schindelin, 2012] [McQuin, 2018] [Berg, 2019]. The main limitation is frequently due to the lack of a flexible 3D viewer to render the resulting processed images (segmented volumes/surfaces) or visualize in an easy and understandable way the overlap between the labels assigned to each object and the original image. Additionally, many functions required for a complete 3D analysis workflow may demand different levels of background knowledge in coding and image analysis.

Considering the growing request for bioimage analysis tools and the difficulties encountered by the users, we developed ZELDA, a plugin for 3D image segmentation and parent-child relation for microscopy image analysis in *napari* [napari contributors, 2019].

ZELDA plugin has the flexibility of being applicable to different purposes and data sets, such as the image measurement of beads to assess microscope resolution (**Fig. 1B**), the RNA quantification in influenza-infected human cell nuclei (**Fig. 2B-D**), the identification of cellular compartments and organelle counting in cell culture samples (**Fig. 4D-F**), or the morphological characterization of organs and tissues (**Fig. 5D-F**).

The 2D and 3D image analysis workflows that ZELDA protocols convey (**Fig. 1A** and **Fig. 2A**) do not require an extensive knowledge of the used algorithms, coding skills, or an elevated number of “point and click” interactions.

The “Data Plotter” protocol (**Fig. 2E**) enables the data exploration during the image analysis, favoring the biological sample comprehension, and potentially highlighting differences between treatments “on the fly”. Furthermore, the reproducibility of workflows is sustained by the implementation of the log (**Suppl. Fig. 2B**) and persistence in memory of the previously used image analysis parameters (i.e. restarting the same protocol will show the parameters values used during the last run).

The implementation of image analysis workflows found in literature is achievable with a fourth protocol called “Design a New Protocol” (**Fig. 3 A-C**). Without any scripting, users can manage the available “widgets” to create a custom GUI (**Fig. 3 E**) that can then be saved and shared with the community (by sharing the JSON database) (**Fig. 3 D**).

Nonetheless, through the customization of the GUI allowed by the fourth protocol, a simply different use of the already available functionalities can lead to better object segmentation. For example, including an additional “Threshold” step after the “Get DistanceMap”, in a newly designed protocol, could help to remove smaller debris before “Show seeds”. Certainly, the possibility of rearranging the components of the image analysis workflows, by using an immediate graphical mode, represents a valuable contribution as an open-source software to bioimage analysis.

To date, ZELDA presents a minimalist interface with three basic protocols implementing image analysis workflows but it could be easily be powered up with additional processing steps to improve image segmentation (e.g. morphological operators to moderate under and over-segmentation, a filter module to exclude segmented objects by intensity or shape descriptors, or allowing to deconvolve the data set before segmenting it).

Although still unable to process images in batch mode, ZELDA can find its niche of application as interactive software since we showed, by benchmarking, that it performs at a comparable level of *ImageJ* and *CellProfiler* in 2D. While in 3D, the segmentation and “parent-child” relation of multi-class objects is performed with a shorter implementation of the workflows and twice faster.

In conclusion, ZELDA plugin for *napari* can hugely accelerate and facilitate the applications of bioimage analysis to life science research.

## 5 Acknowledgements

We thank Sara Barozzi (IEO, Milan) for the preparation of the PML NB sample shown in Suppl. Fig. 1A.

We thank Olivia Swann (Barclay Lab, Imperial College London) for supplying the influenza infected cells analyzed in Suppl. Fig. 1D.

## 6 Conflict of Interest

The authors declare that the research was conducted in the absence of any commercial or financial relationships that could be construed as a potential conflict of interest.

## 7 Authors contribution

RDA developed the *napari-zelda* plugin, acquired and analyzed the data, and wrote the manuscript. GP helped to develop the plugin and wrote the manuscript.

## 8 Funding

RD’A works at the Francis Crick Institute which receives its core funding from Cancer Research UK (FC001999), the UK Medical Research Council (FC001999), and the Wellcome Trust (FC001999).

## 9 Data Availability Statement

The datasets [GENERATED/ANALYZED] for this study can be found in the https://github.com/RoccoDAnt/napari-zelda.

## Footnotes

1 Center for Open Bioimage Analysis: https://openbioimageanalysis.org/

2 Kota Miura 2020, “In Defense of Image Data & Analysis Integrity” - [NEUBIASAcademy@Home] Webinar https://www.youtube.com/watch?v=c_Oi2HKom_Y

## Notes

https://github.com/RoccoDAnt/napari-zelda

### Competing Interest Statement

The authors have declared no competing interest.

