## Supplementary Figures for "ZELDA: a 3D Image Segmentation and Parent-Child relation plugin for microscopy image analysis in *napari*"

### *Supplementary Material*

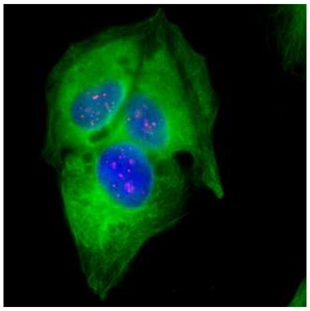

A

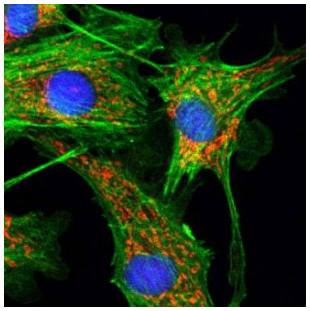

B

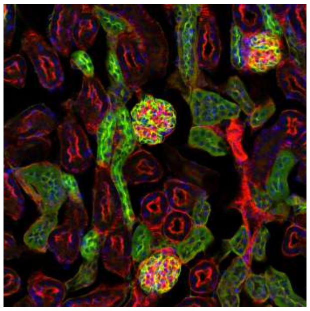

C

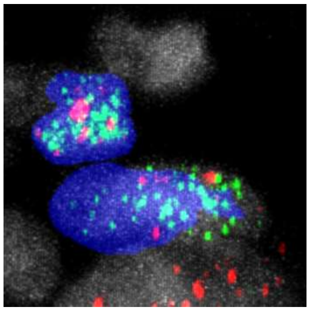

D

**Supplementary Figure 1. Example of biological application of image analysis.** (**A**) Counting of PML NB shells (red) in cell nuclei (blue) is referred to as a “2D counting” task. The cytoplasmic staining is shown in green. (**B**) Segmentation of organelles, such as mitochondria (red), identified as children objects, and their correct assignment to the parent cell cytoplasm (green) as an example of “2D segmentation” and “Parent-child relation”. (**C**) Characterization of structure with WGA and phalloidin staining (green and red) and number of cells with DAPI (blue) in kidney tissue glomeruli is a “3D cell counting” or “3D object segmentation” image analysis task. (**D**) Localization of positive-sense (green) and of negative-sense (red) RNA aggregates within infected cells (blue) or non-infected nuclei (gray) during viral translation or replication in influenza infected human cells, by applying “3D object segmentation” and “Parent-child relation” algorithms.

| | Parent_label | Area | Equivalent_diameter | MFI |
|---|---|---|---|---|
| 0 | 0 | 26 | 3.675571781 | 5628.38462 |
| 1 | 0 | 2 | 1.563185284 | 3498.5 |
| 2 | 0 | 2 | 1.563185284 | 3345.5 |
| 3 | 0 | 3 | 1.789400458 | 3237 |
| 4 | 0 | 3 | 1.789400458 | 3097 |
| 5 | 0 | 6 | 2.254503304 | 4403 |
| 6 | 0 | 8 | 2.481401964 | 4081.5 |
| 7 | 0 | 1 | 1.240700982 | 3113 |
| 8 | 0 | 2 | 1.563185284 | 3181 |
| 9 | 0 | 1 | 1.240700982 | 3088 |
| 10 | 0 | 1 | 1.240700982 | 3028 |

A

```
-> Th=0.5
-> GB: sigma=1.0
-> DistMap
-> Maxima: min_dist=1
-> Found n=7 objects
```

B

**Supplementary Figure 2.** (**A**) Example of exported results table for the "Segment two populations and relate" protocol. The table includes the original label of the children population, the parent identifier, and the measured properties such as the Area, the Equivalent diameter, and the Mean Fluorescence Intensity (MFI). (**B**) A log file is saved as .txt at the end of the segmentation protocols.

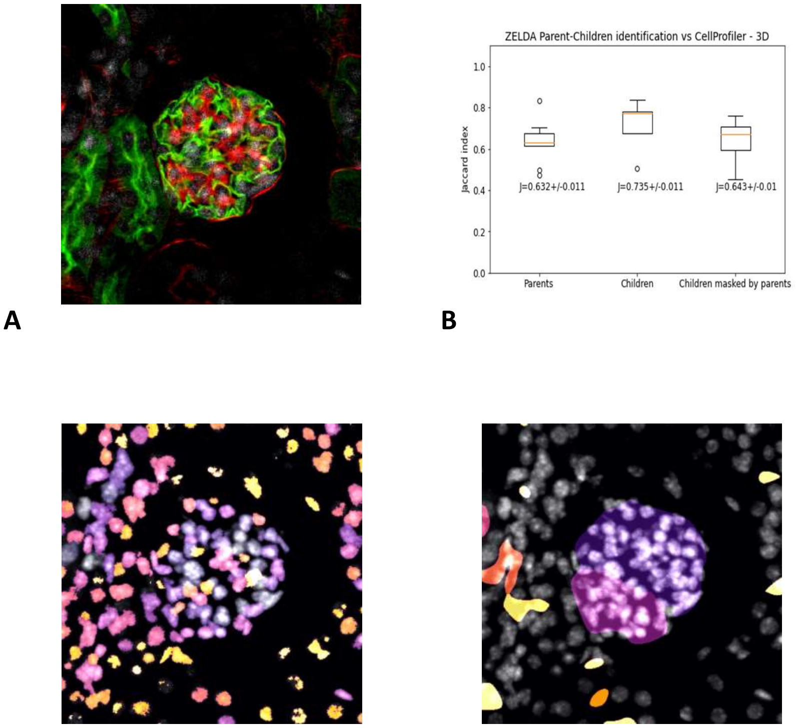


**Supplementary Figure 3.** (**A**) Mid-slice of the z-stack, where typically the strongest signal is found. Its segmentation with ZELDA and *CellProfiler* provide a higher Jaccard score. (**B**) Jaccard index for the results of the ZELDA 3D segmentation and a *CellProfiler* pipeline composed solely of 3D compatible modules. Improved (**C**) segmentation of children nuclei and (**D**) "parent-child" relation obtained by *CellProfiler* using 3D pre-processed data sets from ZELDA.
